# Autophagy in lymphatic endothelial cells promotes lung metastasis

**DOI:** 10.64898/2026.02.22.707064

**Authors:** Diede Houbaert, Kathryn A Jacobs, Patrizia Agostinis

## Abstract

Macroautophagy/autophagy is the main lysosomal pathway for the degradation and recycling of cytoplasmic cargo, with emerging roles in endothelial cell (EC) biology. While autophagy has been extensively studied in blood ECs, its function in lymphatic ECs (LECs) remains unexplored. Given the central role of the lymphatic system in antitumor immunity and metastatic spread, we investigated how LEC autophagy affects metastatic lung colonization. In line with previous reports showing that autophagy regulates the availability of the egress signal sphingosine 1 phosphate (S1P) in secondary lymphoid organs (SLOs), lungs of non-tumor-bearing mice with LEC-specific genetic deletion of *Atg5* (ATG5*LEC-KO* mice) exhibited reduced lymphocyte infiltration. Remarkably, in tumor-bearing mice, either genetic loss of LEC-autophagy or pharmacological blockade of S1P lyase by 4-deoxypyridoxine (DOP) suppressed lung metastasis. Pulmonary immune profiling revealed that while DOP enhanced effector T cell activity despite lower numbers, LEC-autophagy-deficient mice markedly increased the B and CD8 T Cell abundance coupled with profound reduction of VEGFR3 expression in the lung lymphatic vasculature. Together, these findings uncover an autophagy-dependent remodeling of the lymphatic routes and immune niches that fosters metastatic seeding and growth in the lung.

## INTRODUCTION

Autophagy, the major lysosomal pathway for the degradation and recycling of intracellular components, is heightened in cancer cells during metastatic dissemination and promotes cancer cell survival at distant sites^1,2^. However, whether and how autophagy in stromal, non-cancerous cells contributes to metastasis formation is still understudied. Cancer cells can induce the formation of new lymphatic vessels, a process called lymphangiogenesis, which is correlated with LN metastasis and poor prognosis^3^. In a recent study, we unraveled that autophagy in LECs boosts inflammation-driven lymphangiogenesis in the cornea by maintaining LEC identity, through facilitating Prox1-mediated VEGFR3 expression^4^. In addition, tumor-associated lymphangiogenesis is abrogated upon loss of LEC-*Atg5*^5^. In the same study, we showed that LEC-autophagy is a major regulator of lymphocyte trafficking by influencing sphingosine-1-phosphate (S1P) gradients within SLOs^5^. Loss of autophagy in LECs increased intranodal S1P levels, in homeostasis and tumor-bearing mice, thereby impairing the egress signal for (naïve) T cells and leading to the retention of (tumor-specific) lymphocytes in (tumor-draining) lymph nodes (TdLNs). Consequently, preventing efflux of ICB-experienced T and NK cells from the TdLNs to the tumor bed induced resistance to immune checkpoint blockade (ICB) therapy^5^. Notably, these effects were largely phenocopied by the pharmacological inhibitor of the S1P lyase DOP, further illustrating that impairing T cells’ exit from the TdLNs is a persuasive consequence of the loss of Atg5 in lymphatic vessels of the LNs^5^.

Interestingly, a recent in vivo genome-wide screen for microenvironmental regulators of metastatic colonization identified the S1P transporter SPNS2 as the major inducer of metastasis formation out of 810 mutant mouse lines that were tested^6^. Both global and LEC-specific knockout of *Spns2* greatly reduced pulmonary metastasis after tail vein injection, which was attributed to an increased percentage of effector T cells and NK cells in the lungs. SPNS2, one of the two main S1P transporters, is expressed in LECs and is necessary in guiding T cell egress from SLOs^7,8^. Interestingly, treatment with DOP phenocopied the antimetastatic effects of *SPNS2* loss, indicating that changes in the circulation of lymphocytes can also affect their effector state and functionality beyond SLOs. Although the lungs are not considered to be a lymphoid organ, lymphatic vessels in the lungs play an important role in interstitial fluid homeostasis, which impact immune cell trafficking and effector function^9^. Combining these data and our previous study, we hypothesized that the knockout of lymphatic *Atg5* could affect lung metastasis, possibly by controlling the trafficking of lymphocytes and remodeling the local lung immune environment.

## RESULTS

To establish the role of LEC-autophagy in metastatic colonization, we performed experimental metastasis assays in mice with a knockout of *Atg5* in the lymphatic endothelial compartment (ATG5^LEC-KO^), by crossing *Atg5*^fl/fl^ mice with *Prox1*-*cre*^*ERT2*^ mice expressing a tamoxifen-inducible Cre recombinase^4,5^. The effects of the conditional deletion of *Atg5* in these mice on autophagic flux of the CD31+/Lyve1+ lymphatic vessels and on inflammation/tumor driven lymphangiogenesis were characterized in our previous studies^4,5^. In line with this, staining for LYVE1+ lymphatic vessels of the inguinal LN, indicated that compared with wild type (WT) mice, ATG5^LEC-KO^ mice displayed a lack of ATG5 signal and reduced autophagosome-bound LC3 immunoreactivity, pointing to impaired autophagy flux (**Fig. 1A,B**) observed in our previous studies.. As murine metastasis models, we used B16F10 melanoma cells, commonly used to study the formation of lung metastases^10,11^, and B16F10 overexpressing the main lymphangiogenic growth factor VEGFC (B16F10 VEGFC) to potentiate local lymphangiogenesis^5^. Furthermore, we also used MC38 colon carcinoma cells to generalize our findings in different cancer cell types.

**Figure 1.**
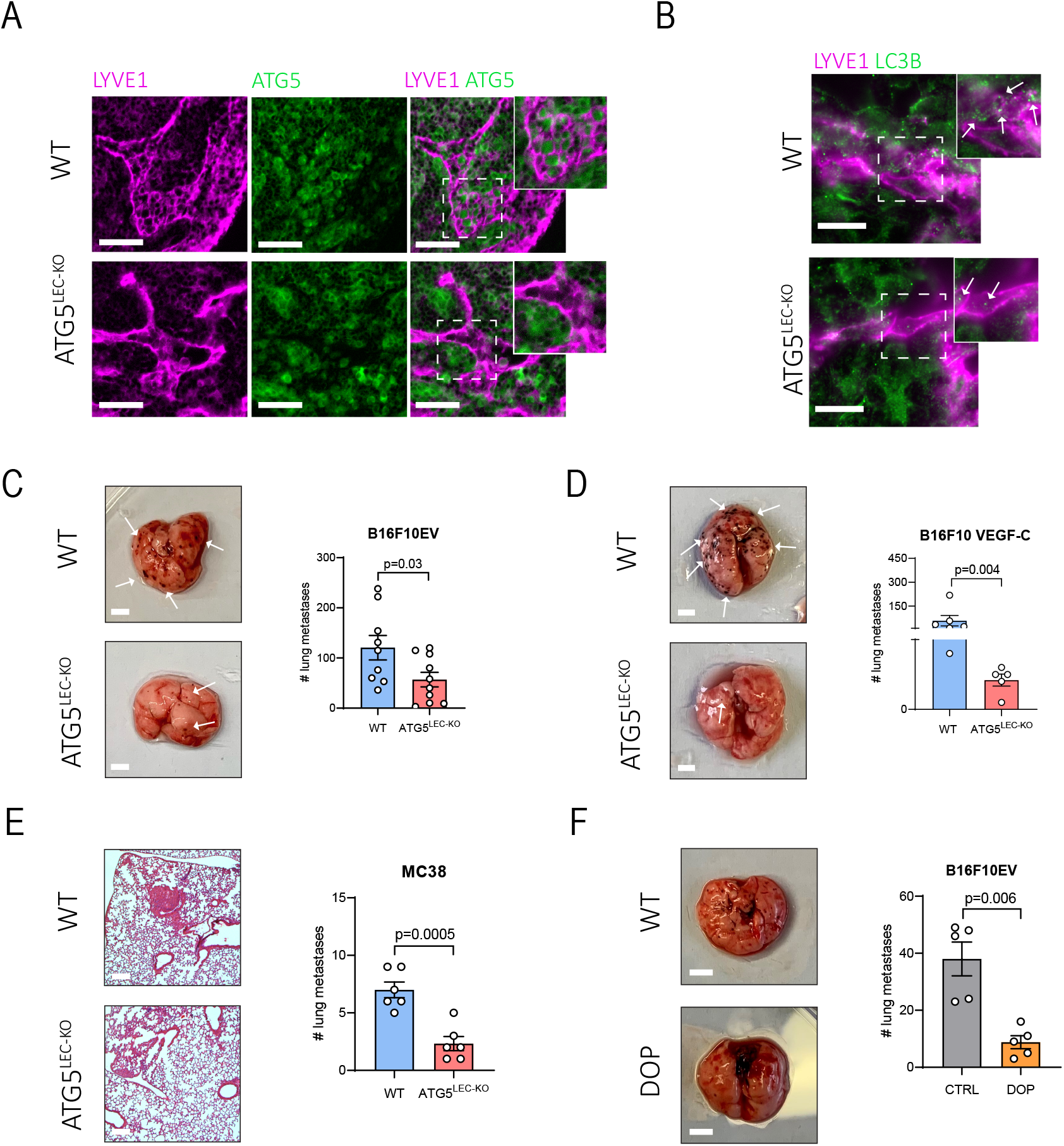
Metastatic colonization is reduced in LEC-specific ATG5 knockout mice. **A)** Representative immunofluorescent images and corresponding crop of LYVE1^+^ lymphatic vessels (magenta) in lymph nodes from wild type (WT) and ATG5^LEC-KO^ mice stained for ATG5 (green) at steady state. A mask following the shape of the lymphatic vessel (LYVE1^+^ staining) was applied (dashed lines). Scale bars represent 50µm. **B)** Representative immunofluorescent images of LYVE1^+^ lymphatic vessels (magenta) in lymph nodes from wild type (WT) and ATG5^LEC-KO^ mice stained for LC3B (green) at steady state. A mask following the shape of the lymphatic vessel (LYVE1+ staining) was applied (dashed lines). Arrows indicate LC3B puncta. Scale bars represent 10µm. **C)** Representative images of metastatic lungs of WT and ATG5^LEC-KO^ mice intravenously injected with B16F10 melanoma cells, 10 days post injection and quantification of lung metastases. Mean ± SEM, n≥9 lungs from independent mice analyzed using unpaired t-test. Scale represents 5mm. **D)** Representative images of metastatic lungs of WT and ATG5^LEC-KO^ mice intravenously injected with B16F10 VEGF-C melanoma cells, 10 days post injection and quantification of lung metastases. Mean ± SEM, n≥9 lungs from independent mice analyzed using Mann-Whitney test. Scale represents 5mm. **E)** Representative images of H&E staining of metastatic lungs of WT and ATG5^LEC-KO^ mice intravenously injected with B16F10 VEGF-C melanoma cells, 10 days post injection and quantification of lung metastases. Mean ± SEM, n=6 lungs from independent mice analyzed using unpaired t-test. Scale represents 1mm. **F)** Representative images of metastatic lungs of CTRL and DOP-treated mice intravenously injected with B16F10 melanoma cells, 10 days post injection and quantification of lung metastases. Mean ± SEM, n=5 lungs from independent mice analyzed using unpaired t-test with Welch’s correction. Scale represents 5mm.

After tail vein injection of B16F10 cells in immunocompetent syngeneic hosts, we found that the number of metastatic nodules was reduced by 50% in ATG5^LEC-KO^ mice compared with their WT counterparts (**Fig. 1C**) and this effect was even more pronounced in mice injected with B16F10-VEGFC melanoma cells with reduction of 90% (n≥5 lungs from independent mice analyzed using unpaired t-test). The pro-metastatic function of LEC autophagy was confirmed using MC38 cells, thus generalizing these findings (**Fig. 1E**). Congruent with a previous report^6^, treatment with DOP reduced metastatic colonization after injection of B16F10 cells (**Fig. 1F**). Together these data show that pharmacological blockade of S1P lyase, recapitulates the antimetastatic effects observed in ATG5^LEC-KO^ mice.

Next, we explored the impact of the loss of LEC-autophagy on the pulmonary immune microenvironment. Following tail-vein injection of B16F10 melanoma cells, we performed immune profiling of lungs isolated from untreated and DOP-treated mice (**Fig. 2A, S1A**). Consistent with its effects on recirculating lymphocytes^6,7^, DOP-treated mice displayed a reduced number of lymphocytes in the lungs as compared with their untreated Remarkably, unlike DOP, WT and ATG5^LEC-KO^ mice did not show consistent alterations in lymphocyte frequency (**Fig. 2B**). On the contrary, mice injected with the more lymphangiogenic cell line B16F10 VEGF-C, exhibited a significant increase in the number of CD8 T cells, B cells and NK cells (**Fig. 2C**). Moreover, when evaluating the spatial localization of B cells (B220), in B16F10-VEGFC model, we noted an increased infiltration throughout the lung tissue of autophagy KO mice compared with WT (**Fig. 2D**).

**Figure 2.**
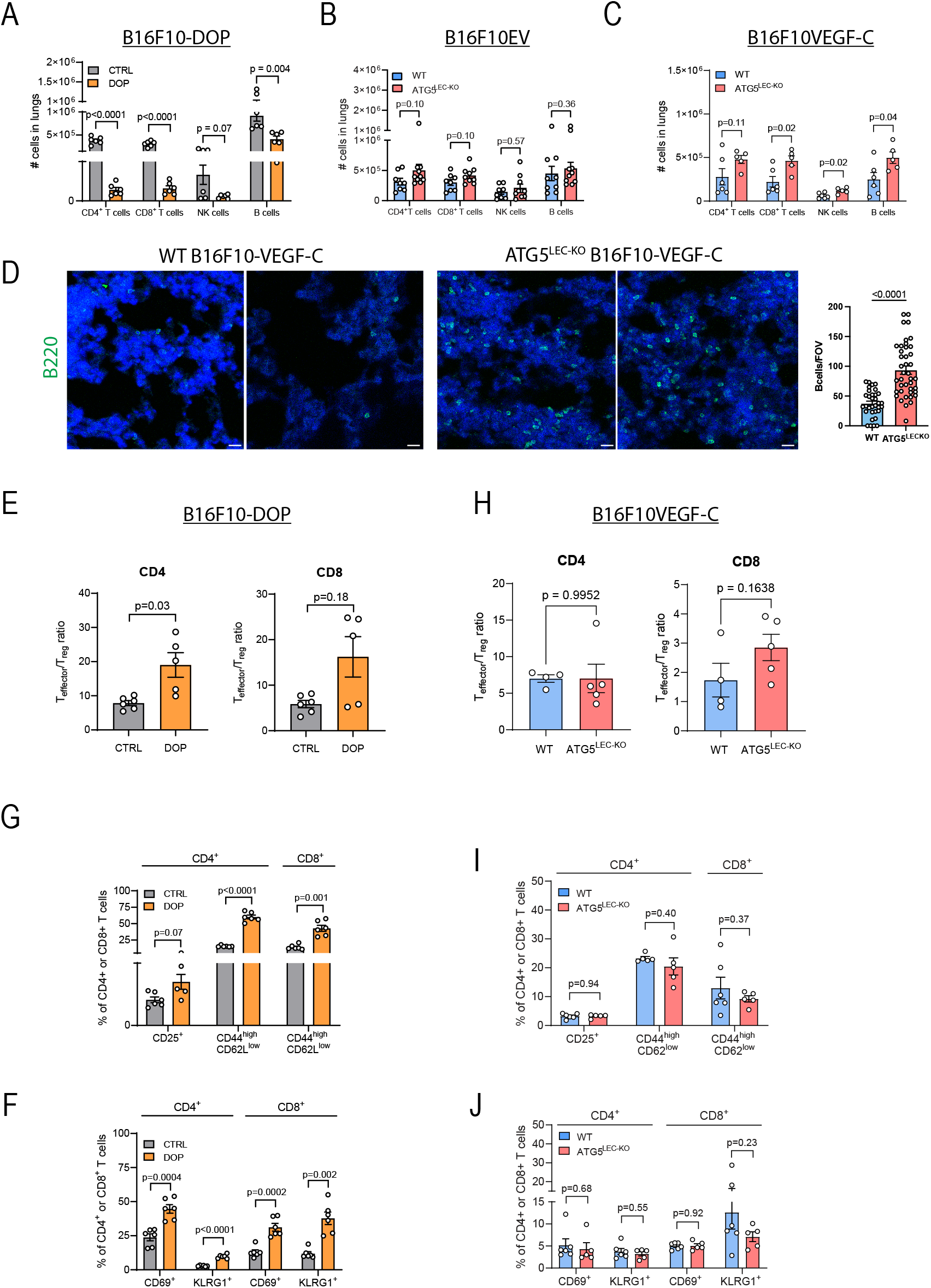
Increased lung immune infiltration in ATG5^LEC-KO^ mice compared to WT mice during metastatic conditions. **A)** Total number of CD4^+^, CD8^+^ T cells (CD3^+^, CD4^+^/CD8^+^), B cells (CD3^-^ CD19^+^) and NK cells (CD3^-^ NK1.1^+^) in lungs of CTRL and DOP-treated mice ten days after intravenous injection of B16F10 cells. Mean ± SEM, n=6 lungs from independent mice analyzed using unpaired t-test (CD4, CD8 and B cells) with Welch’s correction (NK cells). **B)**Total number of CD4^+^, CD8^+^ T cells, B cells and NK cells in lungs of WT and ATG5^LEC-KO^ mice ten days after intravenous injection of B16F10 cells. Mean ± SEM, n≥8 lungs from independent mice analyzed using unpaired t-test (CD8) or Mann-Whitney test (CD4, B, NK cells). **C)**Total number of CD4^+^, CD8^+^ T cells, B cells and NK cells in lungs of WT and ATG5^LEC-KO^ mice ten days after intravenous injection of B16F10-VEGFC cells. Mean ± SEM, n≥5 lungs from independent mice analyzed using unpaired t-test (CD4, B cells) or Mann-Whitney test (CD8, NK cells). **D)** representative immunofluorescence staining and quantification per FOV of B cells (B220) in lung tissue from WT and ATG5^LEC-KO^ B16F10-VEGFC bearing mice, scalebar=20um n≥3 lungs from independent mice analyzed using unpaired t-test **E)** Ratio of CD4 and CD8 effector (CD44^high^ CD62L^low^) T cells on regulatory T cells (CD4^+^ CD25^+^) in lungs of CTRL and DOP-treated mice 10 days after intravenous injection of B16F10 melanoma cells. Mean ± SEM, n≥9 lungs from independent mice analyzed using unpaired t-test with Welch’s correction.**F)** Frequency of Tregs (CD25^+^) and effector (CD44^high^ CD62L^low^) T cells of total CD4^+^ and CD8^+^ T cells in lungs of CTRL and DOP-treated mice ten days after intravenous injection of B16F10 cells. Mean ± SEM, n=6 lungs from independent mice analyzed using unpaired t-test with Welch’s correction. **G)** Frequency of CD69^+^ and KLRG1^+^ T cells of total CD4^+^ and CD8^+^ T cells in lungs of CTRL and DOP-treated mice ten days after intravenous injection of B16F10 cells. Mean ± SEM, n=6 lungs from independent mice analyzed using unpaired t-test (CD69) with Welch’s correction (KLRG1). **H)** Ratio of CD4 and CD8 effector T cells on regulatory T cells in lungs of WT and ATG5^LEC-KO^ mice 10 days after intravenous injection of B16F10 VEGF-C melanoma cells. Mean ± SEM, n≥4 lungs from independent mice analyzed using unpaired t-test (CD8) with Welch’s correction (CD4 **I)** Frequency of Tregs and effector T cells of total CD4^+^ and CD8^+^ T cells in lungs of WT and ATG5^LEC-KO^ mice ten days after intravenous injection of B16F10 VEGF-C cells. Mean ± SEM, n≥5 lungs from independent mice analyzed using unpaired t-test. **J)** Frequency of CD69^+^ and KLRG1^+^ T cells of total CD4^+^ and CD8^+^ T cells in lungs of WT and ATG5^LEC-KO^ mice ten days after intravenous injection of B16F10 VEGF-C cells. Mean ± SEM, n≥5 lungs from independent mice analyzed using unpaired t-test.

Notably, the increase in lymphocyte frequency observed in LEC-*Atg5* KO conditions was specific to metastasis conditions; since in homeostasis lungs of *Atg5*-KO mice displayed a reduced number of lymphocytes as compared to WT lungs (**Fig. S1B**), consistent with the defect in T cell egress from SLOs in steady state conditions described in our previous study^5^.

Furthermore, despite the lymphopenia caused by DOP treatment, lungs of DOP-treated mice displayed a higher Teffector/Treg cell ratio for CD4+ and a trend for CD8 T+ cells (Fig. 2 E,) as well as increased percentage of activated CD44^high^/CD62L^low^ effector CD4+ and CD8+ T cells (Fig. 2F,G,S1C), suggesting that an increased local immunosurveillance could prevent further metastatic colonization under these conditions. Unlike the effects triggered by DOP, ATG5^LEC-KO^ mice bearing B16f10-VEGFC tumors showed no significant changes both in the Teffector/Treg cell ratio and proportion of activated CD44^high^/CD62L^low^ effector CD4+ and CD8+ T cells (Fig. 2H,I,J). Thus, while their antimetastatic effects are similar, the pulmonary immune phenotypes of DOP-treated mice or ATG5^LEC-KO^ mice are pronouncedly different.

Given the discrepancy in phenotype between DOP-treated mice and B16F10-VEGFC, we next probed further into the antimetastatic effects of ATG5^LEC-KO^ mice by evaluating lymphangiogenesis using the B16F10-VEGFC experimental metastasis model. We previously showed that autophagy can affect LEC identity and inflammation-driven lymphangiogenesis by stimulating lipophagy-mediated mitochondrial fatty acid oxidation and the epigenetic control of Prox1 target genes, including Vegfr3^4^. Consistently, immunofluorescence staining of lungs of B16F10-VEGFC-bearing wild-type and ATG5^LEC-KO^ mice revealed that loss of LEC-autophagy resulted in a significantly reduced Vegfr3 vessel density and signal intensity (**Fig. 3A-C**), indicating the inability of lymphatic vessels to grow in response to melanoma-induced lung lymphangiogenesis.

**Figure 3.**
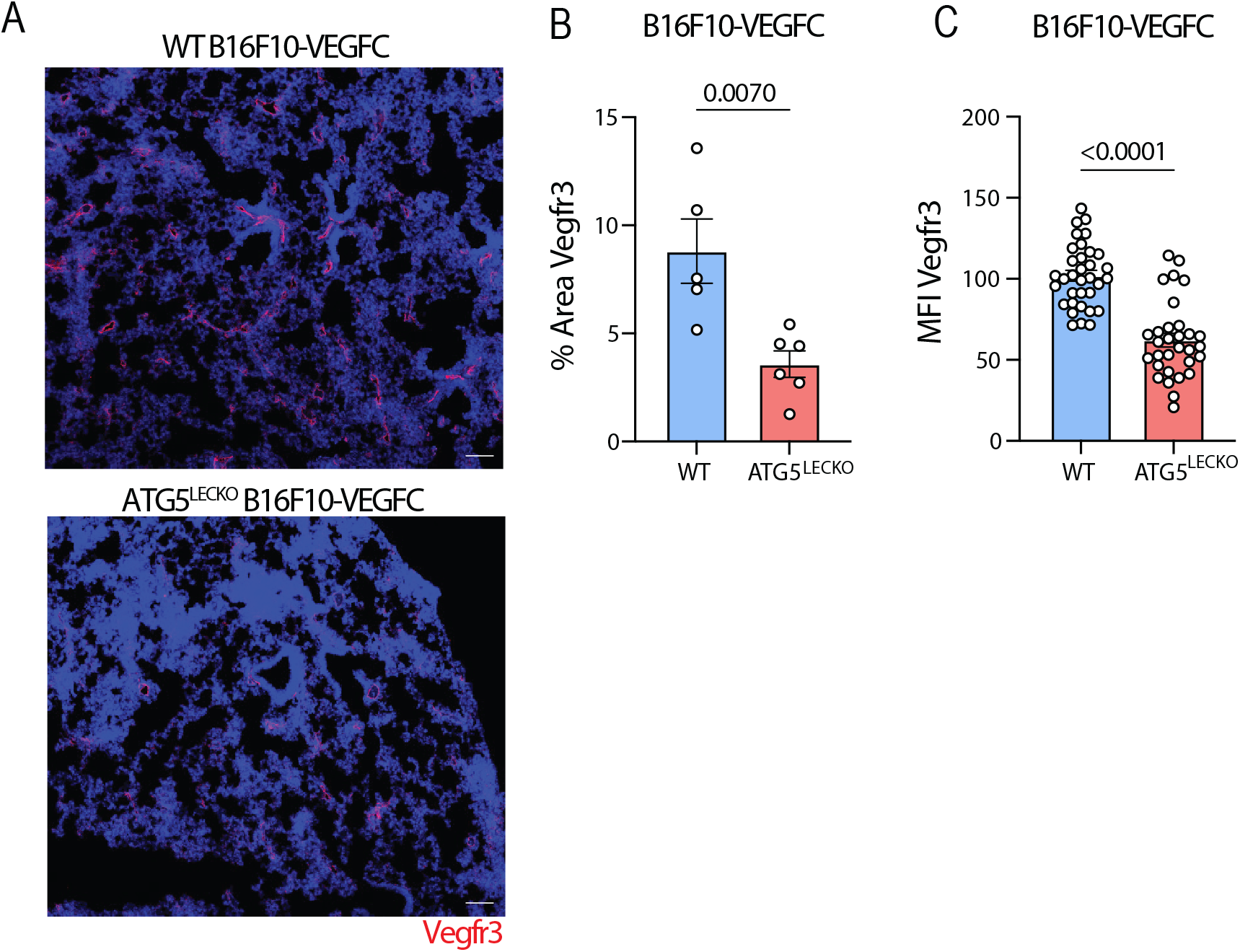
Lymphangiogenesis is blunted in ATG5^LEC-KO^ mice during B16F10-VEGFC experimental metastasis. **A)** Representative immunofluorescence images of Vegfr3 (Red) in WT vs ATG5^LEC-KO^ B16F10-VEGFC metastasis bearing mice. Scalebar=100um. **B)** Quantification of Vegfr3-positive vessel density per lung section. Each dot represents 1 mouse. Mean ± SEM, n≥5 lungs from independent mice analyzed using unpaired t-test **C)** Quantification of mean fluorescence intensity (MFI) of Vegfr3 staining in (**A**). Each dot represents 1 vessel. Mean ± SEM, n≥5 lungs from independent mice analyzed using unpaired t-test

## Discussion

Our report unveils that loss of ATG5 in lymphatic vessels creates a lung microenvironment unfavorable for the growth of metastasizing cancer cells.

Using a mouse model of genetic loss of the key autophagy gene Atg5 in lymphatic endothelial cells, and comparing it with the effects of DOP-mediated inhibition on S1P-lyase on lymphocytes recirculation, we found that lung metastasis was suppressed under both conditions, but through distinct mechanisms. DOP likely exerted its anti-metastatic effects by enhancing lung immunosurveillance, as previously reported^6^. In contrast, loss of Atg5 in LECs impaired tumor-driven lymphangiogenesis and led to a pronounced accumulation of pulmonary B cells and CD8 T cells. Although this immune enrichment may contribute to local antitumor activity, the remarkable suppression of lymphangiogenesis suggests that autophagy deficiency in LECs primarily hinders metastasis by disrupting lymphatic remodeling that supports tumor growth. In line with this, previous studies have shown that enhanced pulmonary lymphangiogenesis, as seen in mice overexpressing VEGF-C in the lung, increases lymphatic vessel density and promotes metastasis in both orthotopic and experimental metastasis models^13^. Furthermore, in melanoma patients with lung metastases, elevated lymphatic density within and around metastatic lesions correlates with poor prognosis^13^.

Although it remains unclear whether tumor-associated lung lymphangiogenesis may enhance lymphatic drainage and immune cell trafficking, it is also plausible that the increased accumulation of immune cells in the lungs of Atg5-LEC-KO mice reflects impaired lymphatic drainage and altered leukocyte traffic during inflammation^14^. Such defects could arise from dysregulated signals governing immune cell recruitment or egress upon loss of autophagy in lymphatic vessels, potentially affecting both fluid clearance and the routing of immune cells to draining lymph nodes. In particular, perturbation of B cell–attracting cues such as CXCL13 or S1P-dependent egress pathways may contribute to the enhanced pulmonary retention or homing of B cells, consistent with their established roles in B cell positioning and trafficking in inflamed tissues^15,16^. We previously demonstrated that inflammation-induced lymphangiogenesis is abrogated in LEC-autophagy-deficient mice, via a lipophagy-mediated mitochondrial-nuclear axis fostering Prox1-driven transcription of key LEC markers, including Vegfr3^4,5^. Moreover, single-cell sequencing analysis of LECs from TdLNs of B16F10 tumor-bearing mice revealed that loss of LEC autophagy selectively impacts LEC subtypes and their likely function, enhancing the retention of lymphocytes in TdLN-without overt changes in lymphangiogenesis in the TdLNs^5^.

Here, we extend these findings showing that in B16F10 VEGF-C metastasis-bearing ATG5^LEC-KO^ mice, metastatic burden is markedly prevented compared with the poorly lymphangiogenic parental B16F10 cell line. Furthermore, Atg5-deficiency in lung LEC blunts tumor-driven lymphangiogenesis and enhances the abundance of B cells and CD8 T cells. Collectively, these observations support a model in which the tissue microenvironment dictates the dominant LEC-autophagy phenotype. Given that Atg5 is deleted pan-tissue in LECs in our model, the profound anti-metastatic effect likely reflects a predominant role for lung-resident lymphatic autophagy under these conditions.

From a translational perspective, the data of this study suggest the use of autophagy inhibitors to prevent metastasis. The use of autophagy modulators in the context of cancer remains however very complicated because of the dual role of autophagy in preventing cancer formation and initiation, while fostering the progression of established cancers and reducing the effects of anticancer therapies^17^. Besides this, autophagy also plays crucial and multifaceted roles in cancer-associated stromal cells, which, depending on the context, may either support tumorigenesis or elicit anti-cancer immune responses^12^. In a parallel study using melanoma bearing mice, we found that autophagy in blood endothelial cells supports the anergic and immunosuppressive function of the tumor vessels, thereby reducing local T cell infiltration and effector function and dampening ICB responses^18^. On the other hand, we showed that deletion of the same essential autophagy gene, *Atg5*, in lymphatic ECs, impairs recirculation of T lymphocytes in steady-state and tumor-bearing mice and blunts ICB antitumor immunity responses. A recent study found that LECs, upon treatment with the chemotherapeutic paclitaxel, upregulate autophagy which increased lymphatic vessel permeability and cancer cell dissemination^19^. These studies thus highlight an unprecedented selective role of autophagy pathways in endothelial cells. The current report further implicates LEC-autophagy in metastasis, pointing to a context-dependent role of autophagy even within the same cell type, which is likely dependent on the tissue microenvironmental niche and stress signals.

Clearly, further research is required to explore the specific role of autophagy in pulmonary lymphatic vessels. Notwithstanding, these studies advocate for the development and selective application of EC-targeted therapies, with the ability to target autophagy in a particular vascular bed^20,21^, in cancer prevention and (combinatorial) therapeutic settings.

## Figure legends

**Supplementary Figure 1:**
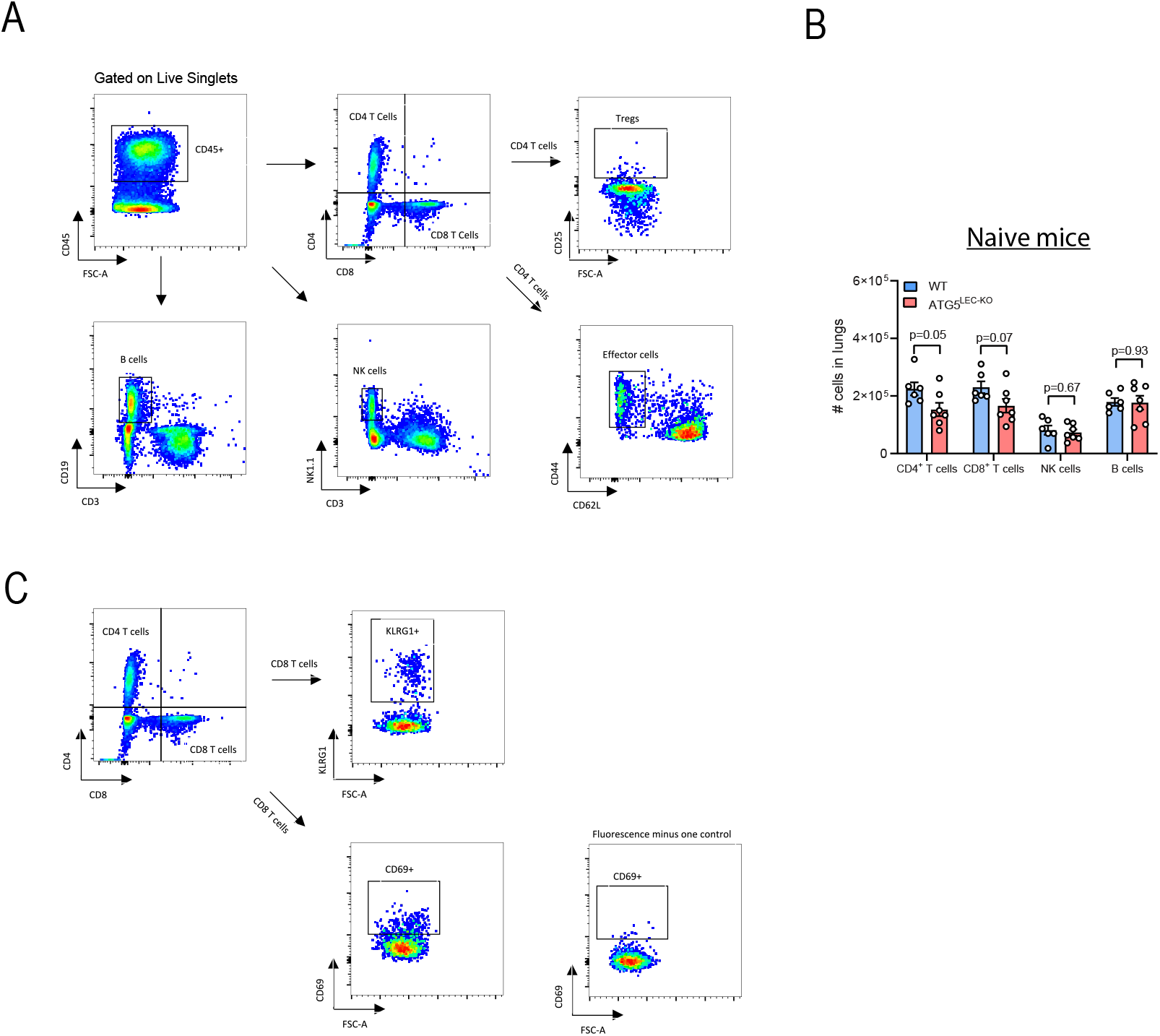
DOP treatment reduces metastasis via increased Teffector/Tregulatory Ratio. **A)** Gating strategies for pulmonary lymphocytes (T cells, B cells and NK cells) and effector and regulatory T cells. **B)** Total number of CD4+, CD8+ T cells (CD3+, CD4+/CD8+), B cells (CD3-CD19+) and NK cells (CD3-NK1.1+) in lungs of WT and ATG5^LEC-KO^ mice at steady state. Mean ± SEM, n≥6 lungs from independent mice analyzed using unpaired t-test. **C)** Gating strategies for CD69+ and KLRG1+ T cells in lungs. Fluorescence minus one control is shown for CD69.

## Materials and Methods

### Mouse models

Animal procedures were approved by the Institutional Animal Care and Research Advisory Committee (KU Leuven) (P167/2018) and were performed in accordance with the institutional and national guidelines and regulations. Mice from the EC-specific inducible Cre-driver line Prox1-cre^*ERT2*^ ^22^ were crossed with Atg5^fl/fl^ mice^23^ to obtain mice with LEC-specific deletion of the *Atg5*gene. These lines were on a 100 % C57BL/6 background. For experiments in this study, we used mice expressing Cre (Prox1-Cre+^*ERT2*^; *Atg5*^*fl/fl*^), referred to as LEC-Atg5^-/-^ and their Cre-negative littermates (Prox1-Cre-^*ERT2*^; *Atg5*^*fl/fl*^), referred to as WT. Tamoxifen (T5648, Sigma Aldrich) injection (i.p.50 mg/kg) was done daily for 5 consequent days 1 week prior to experimental procedures. These lines were on a 100% C57BL/6 background. 4-deoxypyridoxine-HCl (DOP, D0501, Sigma Aldrich) treated mice received drinking water with 10g/L glucose plus 100mg/L DOP, vehicle-treated mice received drinking water with 10g/L glucose.

### Experimental metastasis assay

300.000 B16F10, B16F10VEGF-C or MC38 cancer cells were injected in the tail vein in 100µl PBS. Mice were sacrificed ten days later. Melanoma metastases were counted macroscopically, while lungs with MC38 metastases were fixed overnight in 4% PFA at 4°C, dehydrated and embedded in paraffin. Sections were cut and stained with hematoxylin and eosin for histopathological examination. Images were acquired using a Leica DM5500 upright microscope (KUL-VIB, CCB), 10x magnification. DOP was given to the mice for a total of 21 days.

### Tissue processing

Pool of LNs (inguinal, axillary, brachial and mesenteric) were mechanically disrupted and filtered through 40µm strainers (43-50040-51, Pluriselect) to obtain single cell solutions. Lung tissues were transferred into a gentleMACS C tube (130-093-237, Miltenyi Biotec) containing digestion medium (KnockOut™ DMEM (10829018,Thermo Fisher Scientific), penicillin/streptomycin (P4333, Sigma Aldrich), 2x Antibiotic-Antimycotic (A5955, Sigma Aldrich), 1 mM sodium pyruvate (S8636, Sigma Aldrich), 1x MEM Non-Essential Amino Acids Solution (M7145, Sigma Aldrich) supplemented with 0.1% collagenase II (17101015, Thermo Fisher Scientific), 0.25% collagenase IV (17104019, Thermo Fisher Scientific) and 15 μg/mL DNase (D4527-10KU, Sigma Aldrich) I. Each sample was further dissociated using the gentle MACS dissociator system (Miltenyi Biotec). All cell suspensions were kept in PEB buffer (0.5% BSA and 2 mM EDTA in PBS).

### Flow cytometry

Cells were incubated with FC block (101319, Biolegend) for 10 minutes and then stained with Fixable Viability Dye eFluor™ 780 (65-0865-14, Thermo Fisher Scientific), anti-CD3 (130-116-530, Miltenyi Biotec), anti-CD4 (130-116-546, Miltenyi Biotec), anti-CD8 (565968, BD Biosciences), anti-CD44 (130-116-495, Miltenyi Biotec), anti-CD62L (565261, BD Biosciences), anti-CD45 (748370, BD Biosciences), anti-CD19 (115545, Biolegend), anti-NK1.1 (746876, BD Biosciences), anti-CD25 (740714, BD Biosciences), anti-CD69 (741478, BD Biosciences), anti-KLRG1 (138423, Biolegend), following the FoxP3/Transcription factor staining buffer set (00-5523-00, Thermo Fisher Scientific). Cells were analyzed using a BD FACSSymphony A5 within 24h of staining and data was analyzed using FlowJo software (v10.8.1).

### Immunofluorescence stainings

Lungs were immediately fixed in 4% PFA overnight at 4°C, dehydrated, and embedded in paraffin or sucrose cryo-preserved (20%) and then embedded in OCT and frozen. Immunostainings were performed using an in-house protocol. Antibodies used were goat anti-LYVE1 (R&D systems, AF2125), rabbit anti-ATG5 (12994S, CST), rabbit anti-LC3B (3868S, CST), Vegfr3 (1:100, 14-5988-82, Thermo Fisher Scientific), Rat anti-B220 (1:100, 103204, Biolegend) overnight at 4°C and corresponding secondary antibodies were added for 1 h at 4°C. Slides were mounted in mounting medium (DAKO) and dried overnight. Confocal images were acquired with a Leica sp8x <u>confocal microscope</u> (KUL-VIB CCB), 60X magnification.

### Quantification and statistical analysis

All data are represented as mean ± SEM. Normality of data was checked using Anderson-Darling, D’Agostino & Pearson and Shapiro-Wilk testing. Statistical significance between two groups was determined by standard unpaired t-test with F-testing or one sample t-test. Unless otherwise indicated, statistical significance between multiple groups was determined by one-way ANOVA to ensure comparable variance, then individual comparisons performed by Dunn’s post-hoc test. In case of non-normality, Mann Whitney or Kruskal Wallis tests were performed. Analysis was done in Prism v9.0f, GraphPad.

